# Single cell RNA sequencing reveals shifts in cell maturity and function of endogenous and infiltrating cell types in response to acute intervertebral disc injury

**DOI:** 10.1101/2024.08.10.607363

**Authors:** Sade W. Clayton, Aimy Sebastian, Stephen P. Wilson, Nicholas R. Hum, Remy E. Walk, Garrett W.D. Easson, Rachana Vaidya, Kaitlyn S. Broz, Gabriela G. Loots, Simon Y. Tang

**Affiliations:** Washington University in St. Louis, St. Louis MO; Physical and Life Sciences Directorate, Lawrence Livermore National Laboratory, Livermore CA; Department of Orthopaedic Surgery, University of California Davis Health, Sacramento, CA, United States

**Keywords:** scRNASeq, IVD, MSC, repair, degeneration, intervertebral disc, spine, cartilage

## Abstract

Intervertebral disc (IVD) degeneration contributes to disabling back pain. Degeneration can be initiated by injury, and progressively leads to irreversible cell loss and loss of IVD function. Attempts to restore IVD function through cell replacement therapies have had limited success due to knowledge gaps in the critical cell populations and molecular crosstalk after injury. Here, we used single cell RNA sequencing to identify the transcriptional changes of endogenous cells of the IVD and infiltrating cell populations following IVD injury. Control and Injured coccygeal IVDs were extracted from 12 week old female C57BL/6J mice 7 days post injury and subjected to single-cell resolution transcriptomic sequencing. Clustering, gene ontology, and pseudotime trajectory analyses determined transcriptomic divergences in the cells of the Injured IVD, flow cytometry identified they types of infiltrating immune cells, and immunofluorescence was utilized to define mesenchymal stem cell (MSC) localization. Clustering analysis revealed 11 distinct cell populations that included IVD, immune, vascular cells, and MSCs. Differential gene expression analysis determined that Outer Annulus Fibrosus, Neutrophils, Saa2-High MSCs, Macrophages, and Krt18^+^ Nucleus Pulposus (NP) cells were the major drivers of transcriptomic differences between Control and Injured cells. Gene ontology revealed that the most upregulated biological pathways were angiogenesis and T cell-related while wound healing and ECM regulation categories were downregulated. Pseudotime trajectory analyses revealed that IVD injury directed cells towards increased differentiation in all clusters, except for Krt18^+^ NP cells which remained in a less mature cell state. Saa2-High and Grem1-High MSCs populations drifted towards more differentiated IVD cells profiles with injury and localized distinctly within the IVD. This study revealed novel MSC populations in a heterogeneous landscape of IVD cell populations during injury, and these cells may be leveraged for future IVD repair studies.

**Lay Summary:** The intervertebral disc (IVD) is a spinal joint that accumulates damage with age but has limited tissue repair capabilities. IVD injuries progress into degeneration, and IVD degeneration is a leading cause of lower back pain. Understanding the cell populations that change and respond to injury will uncover targets to restore IVD function. Mesenchymal stem cells (MSCs) are cells within the IVD that can potentially replenish the cells lost after IVD damage. To better understand how IVD cell populations are affected by tissue damage, we performed single cell RNA sequencing of IVD tissue 7 days post injury and analyzed the differences in gene regulation. We identified diverse cells populations such as IVD, immune cells, vascular cells, and MSCs. We discovered the presence of *Saa2* and *Grem1* expressing MSCs that become less stem cell-like and express higher levels of IVD gene markers after injury. We also determined that *Saa2* and *Grem1* have varying gene expression patterns in IVD tissues that becomes attenuated after injury. These MSCs could be targeted for future stem cell therapies to prevent IVD degeneration.

## Introduction

Low back pain is a leading cause of disability worldwide and intervertebral disc (IVD) degeneration is a major contributor to back pain^1,2^. IVD degeneration is the progressive deterioration of IVD structure and function and can be instigated by injury, aging, or mechanical instability^3^. The etiology of IVD degeneration and how endogenous cell populations and infiltrating cells participate in the response to IVD damage is still poorly understood. The IVD has a limited reparative capacity, and the lack of innate mechanisms to stimulate repair and restore tissue homeostasis is likely why the IVD is prone to cumulative degeneration^4,5^. Understanding how IVD cell populations respond to injury and identifying resident stem cell populations can provide cellular targets to increase IVD repair and prevent degeneration.

The IVD is essential for spine function and consists of three major tissue types: the annulus fibrosus (AF), the nucleus pulposus (NP), and the cartilaginous end plate (CEP). The AF can be further divided into the inner (iAF) and outer (oAF) annulus fibrosus. The IVD is an avascular and aneural structure that is immunoprivileged when healthy, but the injured IVD becomes infiltrated with nerves, vasculature, and proinflammatory immune cells^6^. The infiltration of these cells during IVD injury and their role in exacerbating chronic degeneration is well characterized, but how endogenous and infiltrating cells affect IVD homeostasis following acute IVD injury response is less defined^7^. Understanding the cellular cascades during the acute injury response, a critical time window to stimulate repair pathways and restore tissue homeostasis, is key to discovering cellular targets to improving IVD repair post injury^8^.

Single cell RNA sequencing (scRNASeq) is a powerful tool to identify novel cell populations and their transcriptional functions at the single-cell resolution. Previous studies have utilized scRNASeq to identify distinct gene markers for the AF and NP, as well as to identify the gene expression profile of the CEP. scRNASeq has elucidated the heterogenous nature of each IVD compartment by uncovering novel tissue markers for the these diverse cell populations within the endogenous tissues (AF, NP, and CEP, and exogenous tissues such as nerves, vasculature, and other cell types including immune cells and mesenchymal stem cells (MSCs)^9-15^. Though previous studies have identified various IVD MSC populations expressing markers such as *Mcam, Ctsk, Cd44, Lglals3*, and *Krt15*, the role of MSCs during IVD pathogenesis is still unclear.

In this study, we subjected C57BL/6 mice to a severe IVD injury to induce degenerative changes. We collected IVD tissue 7 days post injury and conducted scRNASeq to analyze the changes in IVD cell populations during the acute injury response and identify novel populations that could be targetable to improve IVD repair strategies. We discovered the most upregulated biological processes were related to angiogenesis and T cell regulation while wound healing and extracellular matrix (ECM) regulation were the most downregulated. We discovered the presence of Inflammatory NP-like cells that are similar to but distinct from other NP clusters that expressed elevated proinflammatory cytokines in conjunction with the increased presence of macrophages and neutrophils with injury. We also identified the presence of two MSC clusters, Saa2 and Grem1 High expressing MCSs, that lose stemness and stimulate differentiation into IVD tissues with injury and localize to IVD tissues based on immunofluorescence. The identification of these novel cell populations and the biological functions they stimulate in Injured IVDs provides targetable cell types to mediate the deleterious changes after IVD injury and stimulate repair.

## Results

### scRNASeq identified 11 clusters consisting of IVD tissues, immune cells, MSCs, and vasculature cells

Unsupervised clustering analysis identified 11 cell populations present in both Control and Injured IVD samples (**Figure 1A**). A heat map of the differentially expressed genes enriched in each cluster relative to all other clusters was used to determine cell identities (**Figure 1B**). We identified intervertebral disc (IVD) tissues represented in clusters 1,2,3,4, and 9 which are outer annulus fibrosus (oAF), Cd24^+^ nucleus pulposus (Cd24^+^ NP), inner AF (iAF), Inflammatory NP-Like cells (Inflamm NP), and Krt18^+^ NP cells, respectively (**Figure 1B**, C). Immune cells were present in cluster 5: Neutrophils, and cluster 8: Macrophages. Two MSC populations were identified: Cluster 6: Saa2-High MSC (Saa2 MSC), and cluster 7: Grem1-High MSCs (Grem1 MSC), and vasculature cell types were also identified: cluster 9: Endothelial cells and cluster 11: Pericytes. Of the 11 clusters identified, the majority showed an increase in cell number with injury except for three clusters: oAF, Grem1 MSC, and Krt18^+^ NP where these cell populations were decreased with injury (**Figure 1C**). Gene expression levels of established tissue markers for the cell types identified in each cluster was measured to support our labeling of the distinct clusters. Most IVD gene markers were expressed in multiple IVD tissues; therefore, quantifying levels of gene expression of each marker was the best strategy for identification. *Col1a1* (**Figure 1D**) and *Lum* (**Figure 1E**) are extracellular matrix proteins known to be expressed oAF and iAF, while *Fmod* (**Figure 1F**), *Col2a1*, (**Figure 1G**), and Acan (Figure 1F) are extracellular matrix proteins highly expressed in the iAF and NP^16-18^. Cluster 4 was identified as Inflammatory NP-Like cells because of the expression of NP tissue markers in conjunction with a higher expression of pro-inflammatory cytokines in this cluster in comparison to the other NP clusters (**Figure S1**). Myeloperoxidase, *MPO*, is an enzyme mainly expressed in neutrophils and this gene was only detected in the Neutrophil cluster **(Figure 1I**)^19^. F4/80 (*ADGRE1*), a murine macrophage marker (**Figure 1J**), and Cd115 (CSF1R), a murine monocyte marker (**Figure 1K**), were both expressed in the Macrophage cluster, highlighting the presence of both cell types in cluster 8^20,21^. Flow cytometry analysis of Cd45^+^ immune cells from Control and Injured IVDs at 7 dpi identified neutrophils, macrophages, and monocytes from Cd11b^+^ myeloid cells and supported our scRNASeq immune population findings (Figure S2). The MSC clusters both highly expressed stem cell markers such as *Cd44*, Sca1, and *mKi67* (Figure S3) in addition to expressing elevated levels of Saa2, Serum amyloid A2, in cluster 6 (Figure 1L), and *Grem1*, a BMP antagonist, in cluster 7 (Figure 1M)^22,23^. Established markers for endothelial cells, Cd31 (*Pecam1*) (Figure 1N), and Pericytes, Thy1 (Figure 1O), were also highly expressed in their respective clusters^24,25^.

**Figure 1:**
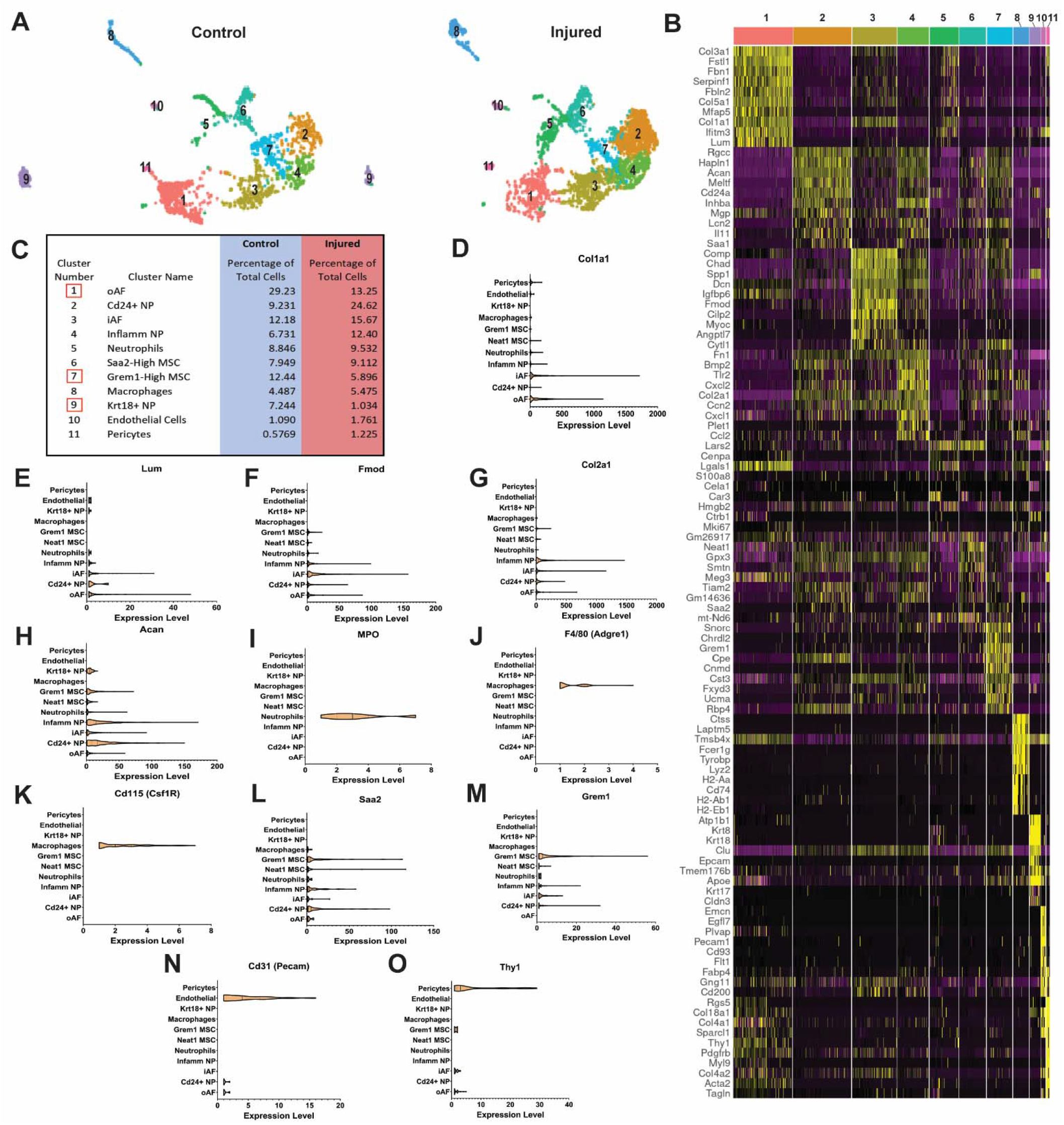
scRNASeq identified 11 clusters consisting of IVD tissues, immune cells, MSCs, and vasculature cell populations. (A) Unsupervised clustering analysis of Control and Injured IVD cells revealed 11 genetically distinct cell populations that include endogenous IVD tissues (AF, NP, and MSC clusters), and infiltrating cell types (immune cells and angiogenic cells). (B) A heat map of the highest expressing genes per cluster aided in the identification of each cluster. (C) The majority of the 11 cell types identified *via* clustering analysis increase in cell number with injury except for 3 clusters: oAF, Grem1 MSC, and Krt18^+^ NP cells (red boxes). The expression of established tissue markers for each cluster was identified to further confirm the presence of oAF with (D) *Col1a1* and (E) *Lum*, iAF/NP with (F) *Fmod*, (G) *Col2a1*, and (H) *Acan*, Neutrophils with (I) MPO, Macrophages and Monocytes with (J) F4/80 and (K) Cd115, respectively, Saa2 and Grem1 MSCs with (L) Saa2 and (M) *Grem1*, Endothelial cells with (N) Cd31, and Pericytes with (O) *Thy1*. IVD- intervertebral disc, AF- annulus fibrosus, NP- nucleus pulposus, MSC-mesenchymal stem cells.

### The acute IVD injury response at 7 dpi revealed increased angiogenesis and T cell recruitment but diminished wound healing and ECM pathways

To assess which RNA transcripts were most regulated due to injury, we analyzed the differentially expressed genes (DEGs) from each cluster and discovered that 295 genes were downregulated, and 122 genes were upregulated in Injured samples relative to Controls (**Figure 2A**). Five clusters, oAF, Neutrophils, Saa2 MSC, Macrophages, and Krt18^+^ NP cells, express the majority of the DEGs (**Figure 2B**). Gene ontology enrichment analysis determined that processes involved in VEGF signaling and T cell regulation were the most enriched from the upregulated DEGs while processes involving extracellular matrix (ECM) catabolism, wound healing, and the inflammatory response are downregulated. The specific genes relevant to each biological process was identified and the clusters from which these genes were differentially expressed are shown to identify the relevant RNA transcripts and cell populations that mediate these biological processes (**Figure 2C**). Neutrophils are tied with Krt18^+^ NP cells for the highest expression of DEGs, and the Neutrophil cluster has the most DEGs implicated in the biological processes identified *via* gene ontology (**Figure 2B,C**). To determine which cell populations were recruiting neutrophils to the IVD after injury, we quantified Cxcl1 expression. Cxcl1 is a potent chemokine that induces neutrophil recruitment and activation in peripheral tissues^26^. We found that *Cxcl1* expression increased with injury in Cd24^+^ NP, iAF, Inflamm NP, Grem1 MSC, and Pericytes (**Figure 2D**). Since biological processes involving T cell regulation were highly enriched from upregulated DEGs, we determined which cell populations regulated *Ccl5* expression, a chemokine that regulates T cell migration^27^. *Ccl5* was only expressed in injured samples and highly regulated in iAF, Inflamm NP, Grem1 MSC and Macrophage clusters (**Figure 2E**). To determine which cell populations were stimulating the pro and anti-inflammatory related biological processes uncovered *via* gene ontology, we measured the expression of proinflammatory cytokines: *IL6* (**Figure 2F**) and TNFα (Figure 2G), and anti-inflammatory cytokines: *IL4* (**Figure 2H**) and *IL10* (**Figure 2I**). IL6 is downregulated in all IVD tissues excepted Cd24^+^ NP and upregulated in MSC and blood vessel specific clusters while TNFα is upregulated majorly in Inflamm NP and macrophage clusters. *IL4* is overall upregulated except for in Cd24^+^NP cells and Krt18^+^ cells which have a downregulation or absence of regulation of *IL4*, respectively. IL10 is upregulated in oAF, Macrophages, Endothelial and Pericyte clusters.

**Figure 2:**
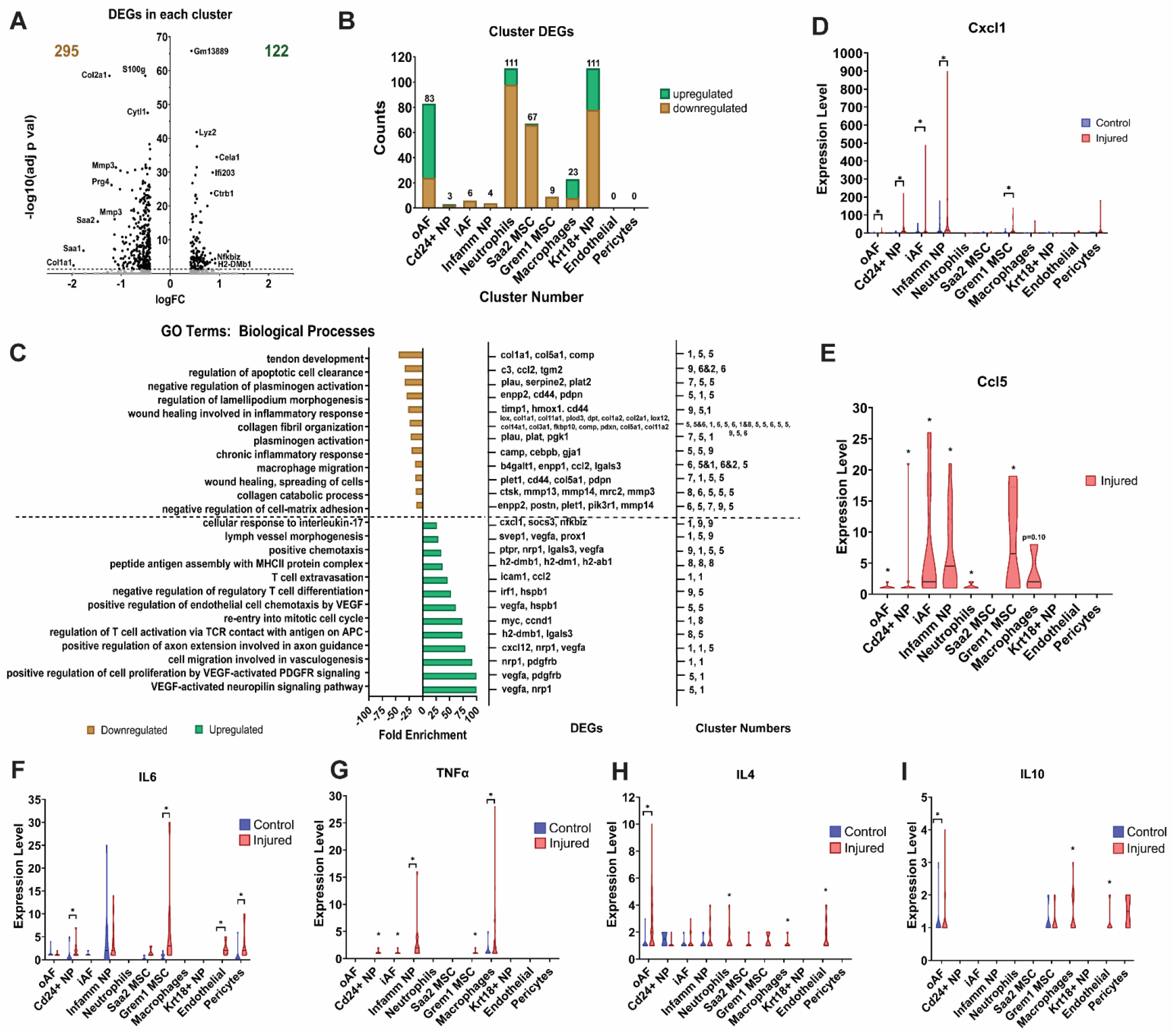
The hallmarks of the IVD injury response at 7 dpi are elevated angiogenesis and increased T cell recruitment but reduced wound healing and ECM pathways. (A) A volcano plot showing the 417 total DEGs from each cluster where 295 are downregulated and 122 are upregulated. (B) The majority of DEGs are expressed in 5 clusters: oAF, Neutrophils, Saa2 MSCs, Macrophages, and Krt18^+^ NP cells. (C) Gene ontology analysis of biological processes revealed the most common upregulated GO terms are angiogenesis and T cell recruitment related while the most common downregulated GO terms are ECM and wound healing related. (D) *Cxcl1* and (E) *Ccl5* are potent regulators of neutrophil and T cell chemotaxis, respectively, and are highly expressed in a cluster specific manner. Proinflammatory cytokines (F) *IL6* and (G) *TNFα* are more highly upregulated with injury than anti-inflammatory cytokines (H) *IL4* and (I) IL10. * = p < 0.05. DEG-differentially expressed genes, GO- gene ontology.

### Injury causes IVD cells to become more terminally differentiated

We performed pseudotime trajectory analysis to determine how the differentiation status of the cell populations changed in response to injury based on differential gene expression for the IVD specific clusters (1-4, and 9) and MSC clusters (6,7). The Monocle package was used to place cells along a pseudotime trajectory corresponding to cell differentiation. Expression data, phenotype data, and feature data were extracted from the Seurat object and a Monocle ‘CellDataSet’ object was constructed using the ‘newCellDataSet’ function. Highly variable genes from Seurat object were used as ordering genes. Trajectory construction was then performed after dimensionality reduction and cell ordering with default parameters. Each cell was sorted in pseudo-timepoints ranging from 0 to 20 where the range indicates an increased change in a cell’s differentiation state or functional state^28^. We discovered a common trajectory path consisting of cells in lower pseudo-timepoints (black arrow) that branched into two distinct cell differentiation paths with higher pseudo-timepoints (red arrows) in both Control (**Figure 3A,C**) and Injured IVDs (**Figure 3B,D**). Analyses of the changes in cell localization on the trajectory branches of each cluster supported a decrease in cells in the lower pseudo-timepoints and an increase in more differentiated cells on the branches with injury (**Figure S4**). Cells from Injured IVDs aggregate at the tail ends of the branches and have higher pseudo-timepoints than when compared to Controls, suggesting increased differentiation of IVD and MSC cells with injury. We identified the IVD cell types that localized on each branch of the pseudotime trajectory by determining where the distinct IVD tissues were concentrated with and without injury (**Figure S4, Figure 3E**). “Fibrocartilaginous-like cells” localized to the top branch since most of the oAF and half of iAF cells were present on this branch. AF cells are described as fibrocartilaginous since they have characteristics of both fibrous and cartilage cells where the oAF is more fibrous-like and the iAF is more cartilage-like and acts as a transition zone between the NP and AF^29^. The bottom branch contains “Chondrocyte-like cells” since half of the cells from the iAF and the majority of the NP cells localized to this branch, and NP cells are described as chondrocyte-like since they share many characteristics with hyaline cartilage^30^ (**Figure S4, Figure 3E**). Interestingly, Krt18^+^ NP cells were the only IVD cell population that were resistant to increased cell differentiation with injury and these cells remained in the lower pseudo-timepoints (**Figure S4**). Quantification of the number of cells present in Controls cells for oAF (Figure 3F), Cd24^+^ NP (**Figure 3G**), iAF (Figure 3H), Inflamm NP (**Figure 3I**), and Krt18^+^ NP (**Figure 3J**) show that the majority are present in low pseudo timepoints (≤ 9)(**Figure 3K**). There was a drastic shift in the differentiation state with injury where the majority of cells in the aforementioned clusters were present in the high (≥ 10) pseudo timepoints except for Krt18^+^ NP cells (**Figure 3L**).

**Figure 3.**
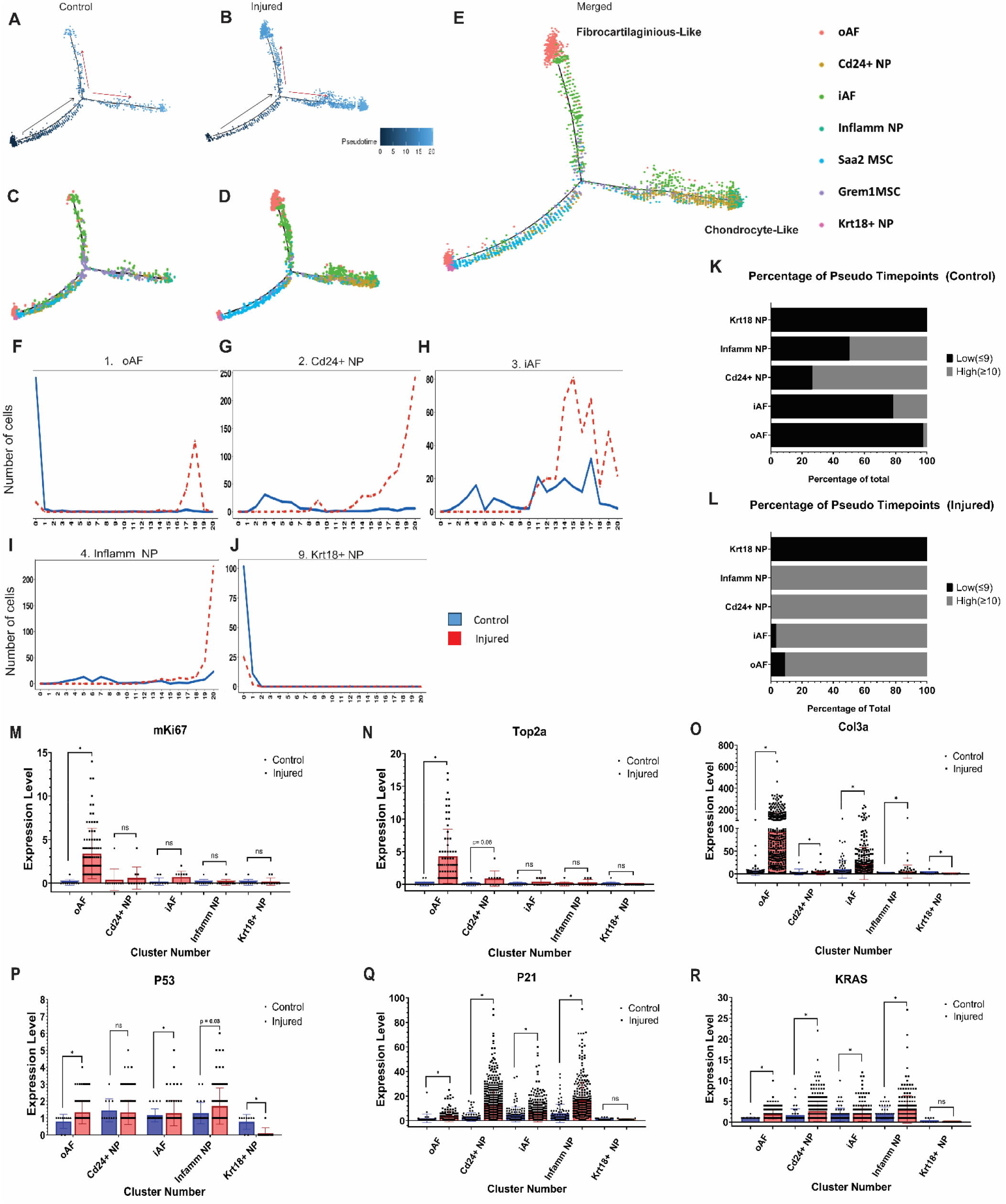
Injury promotes IVD tissues to become more terminally differentiated. Pseudotime trajectory analysis of the (A, C) Control and (B, D) Injured cell clusters from the IVD, 1-4 and 9, show two distinct cell differentiation trajectories where cells are in a more mature cell state in the branches. (E) Identification of the cell type represented on each trajectory branch based off IVD cluster localization. Control and Injured cells from IVD tissue clusters 1-4 and 9 and MSC clusters 6 and 7 were sorted into pseudotime points to identify their cell state maturity where 0 is least mature and 20 is most mature. There is an increase in the number of Injured IVD tissue cells in the more mature cell state pseudotime points in (F) oAF, (G) Cd24^+^ NP, (H) iAF, and (I) Inflamm NP when compared to Controls but not (J) Krt18^+^ NP cells. The percentage of cells from clusters 1-4, and 9 that are in low pseudotime points (≤9) or high pseudotime points (≥10) from (K) Control or (L) Injured samples were quantified. Proliferation and connective tissue healing makers (M) *mKi67*, (N) *Top2a*, and (O) *Col3a* gene expression is regulated in a cluster specific manner but is less regulated than senescence factors (P) *P53*, (Q) *P21*, and (R) *KRAS* with injury. Panels M-R are bar graphs plotted to show the number of individual cells expressing each gene and statistics were ran on gene expression. Mann-Whitney T- tests were conducted to assess statistically significant changes in gene expression between the Control and Injured groups. * = p < 0.05.

Since we observed advanced cell differentiation states based on trajectory analysis and reduced wound healing and collagen fibril organization with gene ontology in Injured IVDs (**Figure 2**), we measured the expression levels of proliferative and connective tissue healing markers with injury. Proliferation markers *mKi67* (**Figure 3L**) and Top2a (**Figure 3M**) were increased in oAF or both oAF and Cd24^+^ NP cells, respectively. *Col3a*, a collagen subtype important for the acute stages of tissue repair, was increased in all 5 IVD tissue specific clusters (**Figure 3O**). We also checked for changes in cell senescence factors with injury to determine if the increase in the number of cells in higher pseudo timepoints correlated with increased cell cycle arrest, a widely reported deleterious consequence of IVD injuries^31^. Senescence-Associated Secretory Phenotype (SASP) factors *p53* (**Figure 3P**), *p21* (Figure 3Q), and *KRAS* (**Figure 3R**) were all upregulated in a cluster specific manner. *P21* and *KRAS* were upregulated in all IVD specific tissue except Krt18^+^ NP cells while p53 was only significantly upregulated in oAF and iAF and downregulated in Krt18^+^ NP cells. These data highlight the cell state changes in IVD tissues in response to injury where increased cell differentiation as measured by pseudo timepoints correlates with increased SASP factor expression and limited regulation of proliferative and repair factors.

### IVD Injury advances cell state maturity and reduces Stem-ness in Saa2 and Grem1 MSC cell clusters

Next, we wanted to assess how the Saa2 MSC and Grem1 MSC populations were responding to injury and if these cell populations were acting as progenitor cells for the AF or NP populations. Pseudotime trajectory analysis determined that cells from the Saa2 MSC cluster were mostly localized to the unbranched region of the pseudotime tree in Controls and there was a slight increase in cell localized to the branches with injury. Grem1 MSC cells were distributed throughout the pseudo timepoints and are on both branches, but the localization becomes more skewed towards the Chondrocyte-like branch with injury (**Figure S4**). These data indicate that Saa2 MSCs largely remain undifferentiated with injury, but cells from Injured samples start to differentiate towards the Fibrocartilaginous and Chondrocyte-like cell fates (**Figure S4F**). However, Grem1 MSCs preferentially undergo differentiation towards the Chondrocyte-like cell fate with injury (Figure S4G). Analysis of the exact pseudo timepoints Saa2 MSCs (**Figure 4A**) and Grem1 MSCs (**Figure 4B**) localize to with and without injury show that Saa2 MSCs are majorly in the low pseudo timepoints in Control and Injured samples (**Figure 4A, C**), but Grem1 MSC cells largely locate to the middle pseudo timepoints (8-15) in Control samples and there is an increase in the highest pseudo timepoints (18-20) with injury (**Figure 4B, D**). These findings support that Saa2 MSCs remain relatively undifferentiated with injury, but Grem1 MSCs from Injured samples become less stem-like and undergo differentiation. To support these observations, we measured the expression of IVD genes markers and highly expressed markers in the IVD clusters to determine if these genes were also increased in the MSC clusters with injury. Saa2 MSC (**Figure 4E**) and Grem1 MSC (**Figure 4F**) clusters both showed an increase in IVD tissue markers throughout the pseudo timepoints with injury relative to Controls. In addition, there was a reduction in stem cell markers in the Saa2 MSC and Grem1 MSC clusters with injury when compared to Controls (**Figure S5**). These data show that the MSC clusters become less stem and more differentiated with elevated IVD tissue marker expression with injury.

**Figure 4.**
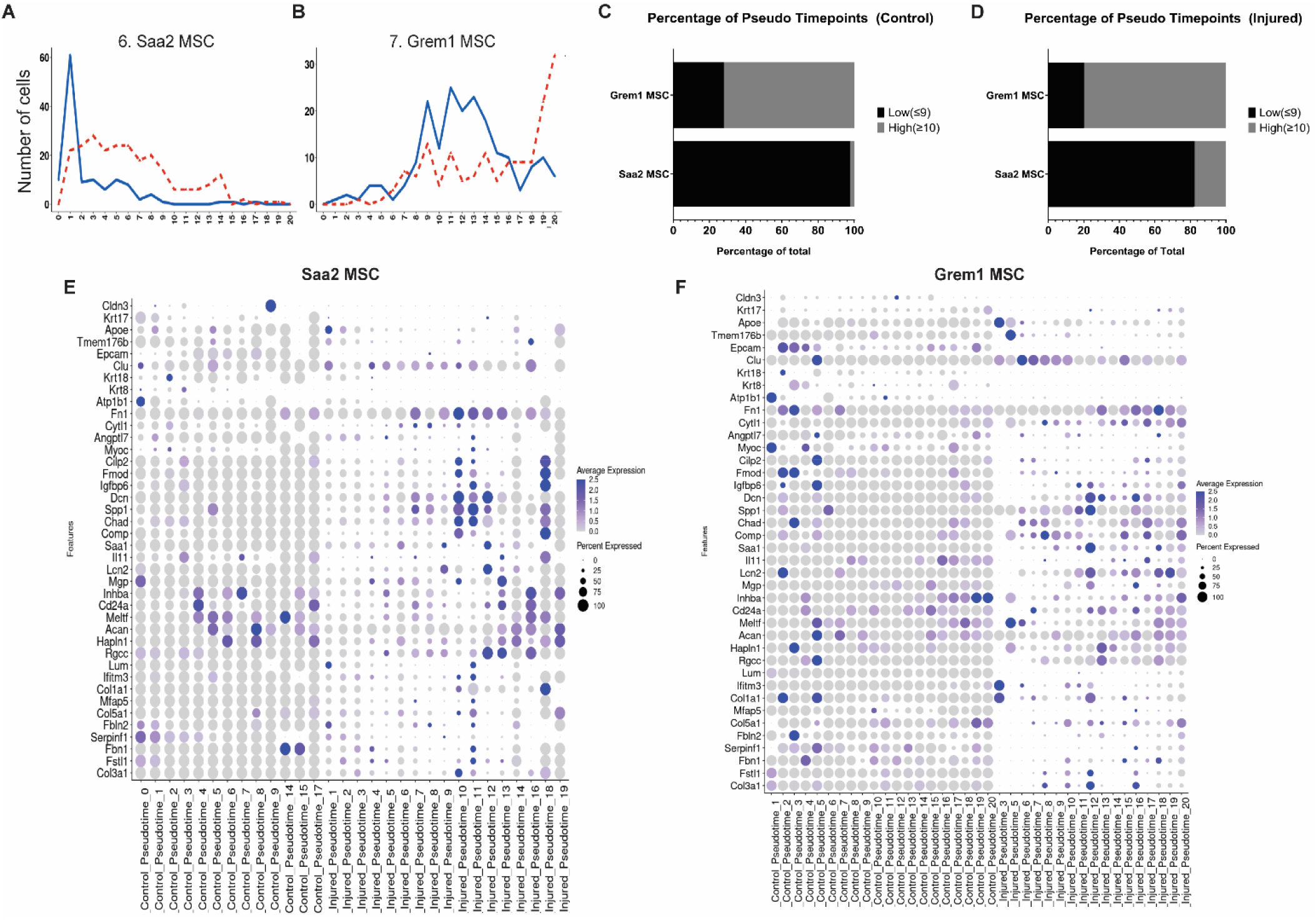
IVD Injury accelerated cell differentiation in Saa2 and Grem1-High MSC cell clusters. (A) Saa2 MSC and (B) Grem1 MSC cells from Injured IVD samples are in more mature pseudotime points than Control cells and the percentage of cells that are in low pseudotime points or high pseudotime points from (C) Control or (D) Injured samples were quantified. Grem1 MSC have a reduced number of cells in a less mature cell state with a sharp increase in cells present in the most mature pseudotime point due to injury while Saa2 MSCs retain more cells in the less mature cell states after injury. Both (E) Saa2 MSC and (F) Grem1 MSCs have an increase in the number and expression levels of IVD tissue markers due to injury.

### Saa2 MSC cells localize to both the Annulus Fibrosus and the Nucleus Pulposus of the IVD

The Saa2 MSC cluster had increased cell numbers and retained stemness with injury based on trajectory analysis (**Figure 1C, Figure S4F**). We utilized immunofluorescence to localize Saa2 positive cells within the IVD and quantify how localization changes with injury. In controls samples, Saa2 localized to the NP, iAF, and vertebrae growth plates (**Figure 6A-A″**). With injury, we observed a reduction of Saa2 staining within the IVD and increased staining within the vertebrae (Figure 6B-B″). Quantification of the overall fluorescence intensity of Saa2 positively stained cells showed a reduction in Injured samples (**Figure 6C**).

### Grem1 MSC cells localize to the IVD and peripheral tissues and are reduced with injury

Grem1 MSC cluster cell number decreased, and the cells become more differentiated towards the chondrocyte-like cell fate with injury (**Figure 1C, Figure S4G**). To confirm the localization of Grem1 expressing cells within the IVD, we utilized immunofluorescence. In controls samples, Grem1 protein localized to the CEP, iAF, the outer regions of the NP closest to the iAF, and the vertebrae end plates (**Figure 6A-A″**). With injury, we observed a reduction of Grem1 staining within the IVD tissues and the cells stained with Grem1 at the endplates appear hypertrophic (**Figure 6B-B″**). Quantification of the overall fluorescence intensity of Grem1 positively stained cells show a reduction in Injured samples, supporting the scRNASeq data (**Figure 6C**).

{Supplemental Figures}

## Discussion

Limited tissue repair is characteristic of the IVD and may lead to deleterious outcomes after damage^5^. Improving the IVD repair capacity by using stem cell therapies has shown promising improvements to IVD health in previous studies but outcomes have a limited or temporary efficacy^32-34^. Understanding how the heterogeneous IVD cell populations interact with resident stems and how this interaction changes with injury will help improve the efficacy of stem cell therapies. scRNASeq is a powerful technique to uncover the transcriptional changes occurring in the IVD in response to injury. This study utilized clustering analysis in conjunction with gene ontology, pseudotime trajectory, flow cytometry, and immunofluorescence to identify established and novel cell populations, quantify transcriptome changes, and determine the effect of injury on resident MSC populations during the acute injury timepoint of 7 dpi. Here, we have identified biological pathways, cell types, and gene markers that are regulated in response to IVD injury with the intent to pinpoint therapeutic targets for future studies that focus on mediating IVD pathology.

Clustering analysis revealed 11 distinct cell populations present in Control and Injured IVDs, including outer and inner AF, Cd24^+^ NP, Krt18^+^ NP, Macrophages, Neutrophils, MSC populations, and vasculature specific cell types, which align with prior scRNASeq studies (**Figure 1A**)^9,10,12,13^. We also discovered novel cell populations, such as Inflamm NP cells, Saa2 MSCs, and Pericytes, and an increase in the number of cells present in Injured samples when compared to Control. Despite the overall increase in cell number with injury, three clusters have a decrease in cell number: oAF, Grem1 MSCs, and Krt18^+^ NP cells (**Figure 1C**). These cell populations may have reduced cell numbers due to being the most sensitive to the increased inflammatory state and loss of tissue homeostasis that occurs acutely post injury in the IVD^7^. Our data supports an increase in inflammation and a decrease in tissue homeostasis by verifying an increase in neutrophils, monocytes, and macrophages *via* flow cytometry (**Figure S2**), showing an increase in proinflammatory gene markers (**Figure S1, Figure 2D-G**), and identifying a decrease in biological pathways specific to wound healing and ECM related processes based on the differentially expressed genes (**Figure 2C**).

Quantification of the number of DEGs expressed by each cluster showed that 5 of the 11 clusters: oAF, Neutrophils, Saa2 MSC, Macrophages, and Krt18^+^ NP, expressed 94% of DEGs with Neutrophils and Krt18^+^ NP cells expressing 53% alone (**Figure 2B**). These data show that these 5 cell types are the drivers of transcriptional changes in response to IVD injury during this acute injury timepoint. Interestingly, there were less upregulated DEGs than those downregulated by injury, but biological pathways enriched by the upregulated DEGs were the most definite and consistent. Gene ontology analysis identified that biological pathways involved in angiogenesis and T cell regulation were the most enriched in upregulated DEGs. Analysis of the specific genes and clusters driving the enrichment of angiogenesis identified pro-angiogenic genes such as *Vegf, Pdgf, Nrp1*, and *Hspb1* regulated primarily by Neutrophils and the oAF (**Figure 2C**). Vegf and Pdgf signaling are well established pro-angiogenic signaling pathways and have been implicated in IVD studies as drivers of increased nerve and vessel infiltration with IVD degeneration^35^. Neuropilins, such as Nrp1, function as coreceptors for Vegf receptors to stimulate vessel growth and maturity^36^. Hsbp1 is released by endothelial cells and cooperate with Vegfa to regulate angiogenesis^37^.

The genes and clusters driving the enrichment of pathways related to T cell regulation involved gene such as *Hs2-dmb1, Iglas3, Irf1, Hsbp1, Icam1*, and Ccl2 which were regulated by Macrophages, Neutrophils, Krt18^+^ NP cells, and the oAF. Hs2-dmb1 plays a role in peptide loading of major histocompatibility complex Class II (MHC II) molecules on antigen presenting cells^38^. Iglas3 is an ECM protein that negatively regulates Cd4 T cell TCR availability^39^. Irf1 is a transcription factor induced by IFNγ signaling and drives differentiation of Cd4 T cells^40^. Icam1 is a co-stimulatory receptor for T cell activation, and Ccl2 is a chemoattractant^41,42^. T cell related pathways are highly enriched based on DEGs at 7 dpi, but intriguingly, no T cells were identified *via* cluster analysis. We have previously profiled the full acute IVD injury response timeline and found that Vegfa, Pdgfa, and T cells genes are not highly regulated at 7 dpi, and there is no discernable increase in regulation of these genes until 10 dpi, 14 dpi, and 17 dpi, respectively^7^. However, the findings from this study show that injured IVDs are primed to increase expression of biological functions related to these genes in a tissue specific manner. Other GO pathways associated with immune cell chemotaxis and infiltration of peripheral tissues such as “cellular response to interleukin-17”, “lymph vessel morphogenesis”, and “positive chemotaxis” were upregulated as well (**Figure 2c**).

Injury also increases the cell maturation state of the cells from IVD tissues. Pseudotime trajectory analysis found an increase in the number of cells present in the later pseudotime points with injury, which represents cells being in an increased maturation or differentiation state, for oAF, Cd24^+^ NP, iAF, and Inflamm NP cells (**Figure 3K,L**)^28^. Interestingly, Krt18^+^ NP cells were resistant to injury-mediated changes in differentiation states and Injured and Control cells remained in the earlier pseudotime points even though the cell number is decreased with injury (**Figure 1C, 3K,L, Figure S4**). Krt18^+^ NP cells also are tied with the Neutrophils cluster for expressing the most DEGs (**Figure 2B**). Krt18 has been suggested to be a stem cell marker for NP cells^43^. These data suggest Krt18^+^ cells could be a promising additional MSC population to target for IVD repair since they retain stemness but are still highly transcriptionally active with injury. The increase in differentiation state of the majority of the IVD tissue is correlated with elevated gene expression of SASP factors with no change in proliferative factors except for the oAF cells (**Figure 3M-R**).

Saa2 MSC and Grem1 MSC populations have slight increases the cell maturation state with Injury as well. The majority of Saa2 MSC cells remain in the lower pseudotime points even with injury, but cells from Injured samples start to differentiate towards the Fibrocartilaginous and Chondrocyte-like cell fates based on their localization on the pseudotime tree branches (**Figure 3E, 4C,D, S4F**). Saa2 MSC cells increase with injury, indicating this population is a great target for IVD regenerative therapies since they are replenished and retain some stemness even with injury, and have the capacity to differentiate into different cell fates. Serum amyloid A (Saa) genes, such as *Saa1*, have been shown to have higher expression in non-degenerate IVD samples from human NP cells when compared to the AF and are expressed in osteoblast MSCs, but most associated with increased expression during the acute phase of inflammation^10,22^. Conversely, *Grem1* is an established skeletal stem cell marker and is expressed in AF progenitor cells^23,44^. We observed that Grem1 MSCs preferentially undergo differentiation towards the Chondrocyte-like cell fate with injury (**Figure S4G**). These data suggest Grem1 MSCs may differentiate towards NP and iAF cells with injury, which have chondrocyte-like cell characteristics. Both MSC populations have a reduction in stem cell markers expression with an increase in IVD tissue markers, further supporting their capacity to utilized as target for future regenerative studies (**Figure 4E,F, Figure S5**). Saa2 and Grem1 both localize to IVD cells based on immunostaining but have slightly different localization patterns. Both Saa2 and Grem1 is present in the NP and iAF while Grem1 also localizes to the end plates and expression is reduced with injury (**Figure 5, Figure 6**). Despite the potential of these MSCs as targetable cell populations in future studies, Saa2 and Grem1 may simply be good markers for these populations but not necessarily targetable genes for modulation of the function of these cells during injury. Other highly regulated DEGS from these clusters should also be considered, such as *Neat1, Gpx3*, and *Meg3* from the Saa2 cluster and *Cnmd, Chrdl2*, and *Ucma* from the Grem1 cluster, all of which have been identified as stem cell or proliferative markers^45-50^. Additional investigations into their functions and localization in the IVD can provide more targets for repair. Another limitation of this study is that IVD cells were flash frozen once rendered into a single cell suspension before being analyzed for scRNASeq. Freezing the cells could have caused additional stress and increased cell loss of more sensitive cell populations. Despite this limitation, our cell population findings are similar to those reported in previous studies^9,12,13^.

**Figure 5:**
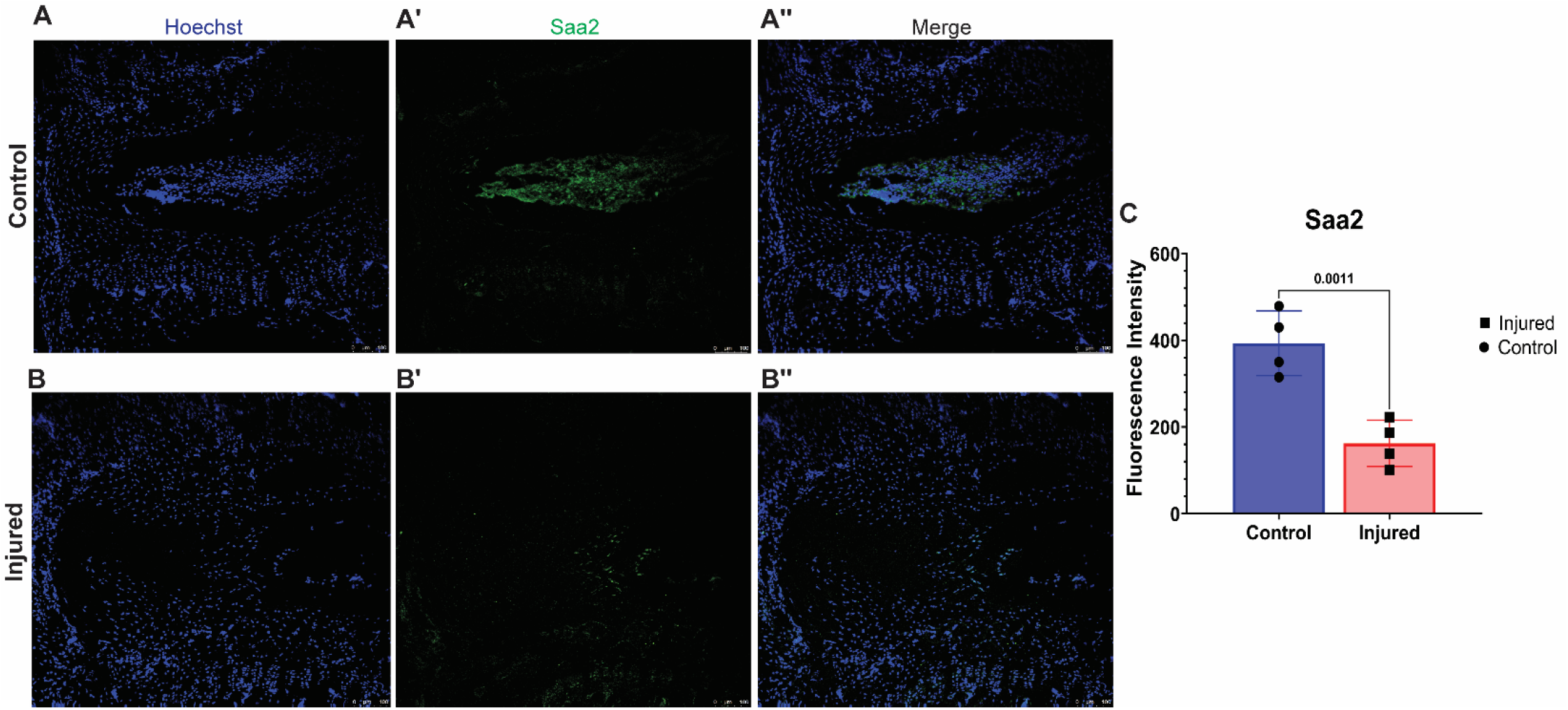
Saa2 positive cells are localized preferentially to the Nucleus Pulposus of the IVD. Saa2 positive cells localize to the NP, inner AF, and peripherally in the vertebrae. There are fewer immuno-stained Saa2 positive cells with injury when comparing (A) Control and (B) Injured IVDs. (C) Quantification of Saa2 immunostaining (n=4).

**Figure 6:**
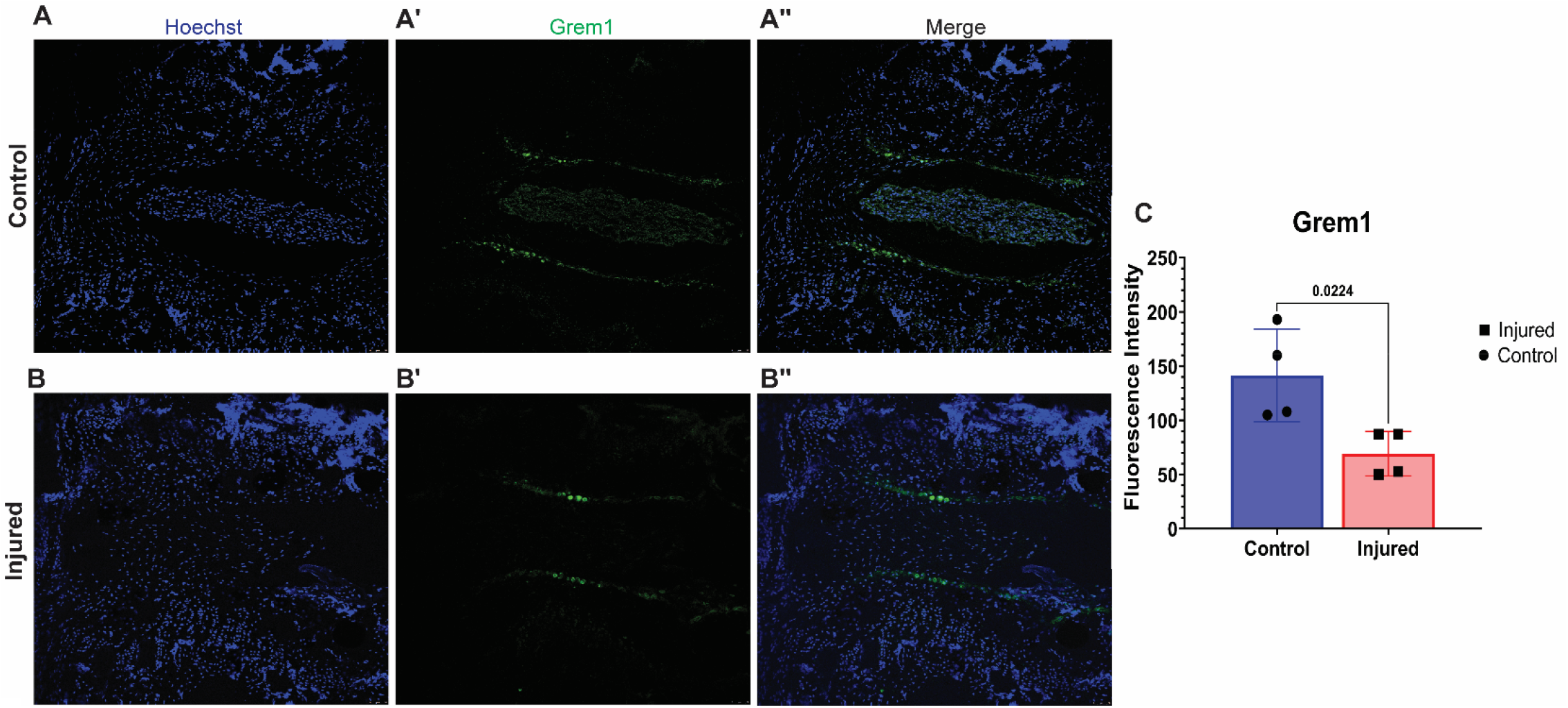
Grem1 positive cells localized to the IVD and bony end plates. Grem1 positive cells localize to the NP, inner AF, and end plates of the IVD, and peripherally in the vertebrae. There is a reduction of Grem1 positive cells when comparing (A) Control and (B) Injured IVDs. (C) Quantification of Grem1 immunostaining (n=4).

In summary, we identified 5 cell populations (oAF, Neutrophils, Saa2 MSCs, Macrophages, and Krt18+ NP) that drive the majority of the transcriptional response to injury at 7 dpi and the biological pathways most regulated due to differential gene expression. We also identified novel populations, such as Inflammatory NP-like cells, Saa2 MSCs, and Pericytes, the role of the Saa2 and Grem1 MSC populations to differentiate into IVD tissues with injury, and the potential of Krt18^+^ NP cells as an additional MSC population. The identification of these novel cell populations and the biological functions they stimulate in injured IVDs provides targetable cell types to mediate the deleterious changes after IVD injury and stimulate repair.

## Methods

### Animal Care

12-week old female C57BL/6J (#000664) mice were purchased from the Jackson Laboratories and housed in a mouse facility under standard laboratory conditions. A chow diet and water were available at libitum. Institutional Animal Care and Use Committee protocols were established at Washington University in Saint Louis and approved before animal usage and conformed to the National Institutes of Health Guide for the care and use of laboratory animals.

### Modeling Intervertebral Disc Degeneration through Injury

Mouse tail IVD injuries were conducted as previously described by our laboratory^35,51^. Control IVDs were isolated from mouse tails in the coccygeal “CC” region CC12/13 through CC16/17. The IVD injury was induced using a 30G needle to bilaterally puncture through the center of the IVD and through both sides of the AF in levels CC5/6 through CC9/10. Control and Injured IVDs are present in each animal. The punctures were performed by using palpation to identify CC5/6 through CC9/10, puncturing each level with a sterile 30G needle, and confirming the accuracy of each puncture with X-ray (Faxitron UltraFocus 100, Hologic). Mice were anesthetized with vaporized isoflurane/oxygen during the procedure and given a single injection of 1mg/mL carprofen at 5mg/kg/mouse as analgesia immediately after injuries. Mice were returned to the mouse facility and monitored every 24 hours until 7 days post injury. For the IVD extractions, mice were euthanized at 7 days post injury (dpi) in a CO2 chamber with 3% CO2 for 5 minutes and a 2 minute dwell time. Euthanized mice were submerged in 70% ethanol for 2 minutes before the IVDs were extracted. IVDs were either rendered into a single cell suspension and counted for single cell RNA sequencing or flow cytometry or intact tissue was processed for immunofluorescence.

### Single Cell RNA Sequencing (scRNASeq)

Control and Injured IVDs from 6 mice were pooled for a total of 30 IVDs in each group. The IVDs were rendered into a single cell suspension by digesting the pooled samples by using four serial digestions of 2mg/mL of Collagenase Type II (Gibco) in DMEM at 37 °C on a rocker for 30 minutes each. The single cell suspension was cryopreserved in a freezing medium 10% DMSO and 90% FBS and snap frozen in liquid nitrogen for storage and shipment. Upon thawing and resuspension in PBS + 0.04% nonacetylated BSA, the single cell sequencing library preparation was performed using a Chromium Single Cell 3⍰ GEM, Library & Gel Bead Kit v3 and Chromium instrument (10X Genomics) following the manufacturer’s protocol, and then sequenced using Illumina NextSeq 500. Alignment of scRNASeq data to the mouse genome (mm10) and gene counting was completed utilizing the 10XGenomics Cell Ranger. Subsequently, output files from the Cell Ranger ‘count’ were read into Seurat v3 for further analysis^52^. Cells with fewer than 250 detected genes or genes that were expressed by fewer than 5 cells were excluded from the analysis. After normalization of the data and the most variable genes identified, the data were scaled, and the dimensionality of the data was reduced by principal component analysis (PCA). A non-linear dimensional reduction was then performed *via* uniform manifold approximation and projection (UMAP) and various cell clusters were identified. All clustering, visualization, and differential gene analysis was performed in Seurat. **Table 1** contains the full list of differentially expressed genes.

### Gene Ontology

Biological pathways enriched by the up-regulated or down-regulated DEGs were identified by using the statistical overrepresentation test from the Panther Classification System with version Panther 18.0. Only GO terms that had a p value and False detection Rate of less than 0.05 were considered. The full list of all GO terms from upregulated DEGS is in **Table 2** and for downregulated DEGs is in **Table 3**.

### Flow Cytometry

Pooled IVDs (n=5 IVDs for each treatment group; n=3 mice for a total of 15 IVDs/flow cytometry run) were isolated and rendered into a single cell suspension by using four serial digestions of 2mg/mL Collagenase Type II (Gibco) at 37 degrees Celsius for 30 minutes each. After resuspension in FACS buffer (0.5% BSA, 2mM sodium azide, 2mM EDTA in 1X PBS), cells were counted and then treated with an Fc blocking buffer containing anti-Cd16/32 (BD Biosciences) for 20 minutes before a 30 minute incubation with antibodies at 4 °C. The antibodies used are as follows: Cd45-APC Cy7 (BD Biosciences), Cd11b-BV605 (Biolegend), Ly6G-BV510 (Biolegend), Ly6C-PE (Biolegend), and 7AAD (Thermofisher) as a live dead stain. For compensation, the spleen from one mouse was rendered into a single cell suspension, treated with Red Blood Cell Lysing Buffer Hybri-Max™ (Sigma Aldrich) and aliquoted to be treated with individual antibodies to serve as single stained controls. After staining, cells were identified by using a LSRFortessa (BD Biosciences) analyzer and FloJo 10.0 software.

### Immunofluorescence

4 IVDs from each group was used for immunofluorescence. 10μm thick, midline, sagittal frozen sections were fixed with 4% PFA for 10 minutes and then permeabilized with 0.5% Triton X in TBS for 10 minutes. Sections were washed in 1x TBS and then blocked using 1% Goat serum in TBS for one hour at room temperature before adding primary antibodies to detect Saa2 (Proteintech, 1:100) or Grem1 (Thermofisher, 1:100) overnight at 4 degrees Celsius. An anti-rabbit Alexa 488 secondary (Thermofisher, 1:250) was added for one hour at room temperature. Slides were stained with Hoechst 33258 (Invitrogen, 1:1000) before being mounted and imaged with a Leica Di8 laser scanning confocal microscope with a 10x objective. Fluorescence was quantified using the analyze particles function on FIJI/Image J.

### Statistical Analyses

Statistical analyses for differentially expressed genes were performed using R statistical software. DEGs from each cluster were identified by having an adjusted p value and false detection rate of p > 0.05. All other statistical analyses were performed using GraphPad Prism version 10.2.0. The assumptions for parametric tests were checked to verify none were violated before statistical tests were ran and results interpreted. For immunofluorescence, differences between Control and Injured samples immuno-stained for Grem1 and Saa2 were analyzed using a paired Student’s T test. A p value < 0.05 was considered statistically significant and an asterisk denotes significance. The magnitude of gene expression for cell-specific markers were tested for differences between Control and Injured cell populations by a Student’s t-test or a Mann–Whitney test as appropriate. Chi-squared tests were used to compare the proportions between matched cell clusters between Control and Injured populations.

## Acknowledgements

This work was conducted with funding support from National Institute of Arthritis and Musculoskeletal and Skin Diseases: R01AR074441, R01AR077678, R21AR081517, P30 AR074992, T32 Postdoctoral Training in Regenerative Medicine (T32 EB028092) from the National Institute of Biomedical Imaging and Bioengineering, the Rita Levi-Montalcini Postdoctoral Fellowship in Regenerative Medicine from the Center of Regenerative Medicine at Washington University, and the Burrough’s Wellcome Fund Postdoctoral Diversity Enrichment Program. Thank you to the Flow Cytometry & Fluorescence Activated Cell Sorting Core for the usage of the core’s analyzers. Part of this work was performed under the auspices of the U.S. Department of Energy by Lawrence Livermore National Laboratory under Contract DE-AC52-07NA27344. This work is preprinted in BioRxiv: doi: https://doi.org/10.1101/2024.08.10.607363.

## Author Contributions

SWC, GGL, and SYT designed the research study, contributed to data interpretation, and wrote the manuscript. AS, SPW, NRH, and GGL performed the scRNASeq analyses. SWC, REW, GWDE, RV, and KSB collected the tissues, performed the research, and analyzed data. All co-authors reviewed and revised the manuscript.

## Data Availability

The raw data files are available on the Gene Expression Omnibus database: GSE277892.

## References

1. Ying Y, Cai K, Cai X, et al. Recent advances in the repair of degenerative intervertebral disc for preclinical applications. Front Bioeng Biotechnol. 2023;11:1259731. doi:10.3389/fbioe.2023.1259731

2. Andersson GBJ. Epidemiological features of chronic low-back pain. The Lancet. 1999/08/14/ 1999;354(9178):581–585. doi:10.1016/S0140-6736(99)01312-4

3. Casiano VE, Sarwan G, Dydyk AM, Varacallo M. Back Pain. StatPearls. StatPearls Publishing Copyright © 2024, StatPearls Publishing LLC.; 2024.

4. Tang G, Zhou B, Li F, et al. Advances of Naturally Derived and Synthetic Hydrogels for Intervertebral Disk Regeneration. Front Bioeng Biotechnol. 2020;8:745. doi:10.3389/fbioe.2020.00745

5. Ju DG, Kanim LE, Bae HW. Intervertebral Disc Repair: Current Concepts. Global Spine J. Apr 2020;10(2 Suppl):130s–136s. doi:10.1177/2192568219872460

6. Lyu FJ, Cui H, Pan H, et al. Painful intervertebral disc degeneration and inflammation: from laboratory evidence to clinical interventions. Bone Res. Jan 29 2021;9(1):7. doi:10.1038/s41413-020-00125-x

7. Clayton SW, Walk RE, Mpofu L, Easson GW, Tang SY. Analysis of Infiltrating Immune Cells Following Intervertebral Disc Injury Reveals Recruitment of Gamma-Delta (γδ) T cells in Female Mice. bioRxiv. 2024:2024.03.01.582950. doi:10.1101/2024.03.01.582950

8. Schultz GS, Chin GA, Moldawer L, Diegelmann RF. Principles of Wound Healing. In: Fitridge R, Thompson M, eds. Mechanisms of Vascular Disease: A Reference Book for Vascular Specialists. University of Adelaide Press © The Contributors 2011.; 2011.

9. Rohanifar M, Clayton SW, Easson GWD, et al. Single Cell RNA-Sequence Analyses Reveal Uniquely Expressed Genes and Heterogeneous Immune Cell Involvement in the Rat Model of Intervertebral Disc Degeneration. Applied Sciences. 2022;12(16):8244.

10. Fernandes LM, Khan NM, Trochez CM, et al. Single-cell RNA-seq identifies unique transcriptional landscapes of human nucleus pulposus and annulus fibrosus cells. Scientific reports. Sep 17 2020;10(1):15263. doi:10.1038/s41598-020-72261-7

11. Gan Y, He J, Zhu J, et al. Spatially defined single-cell transcriptional profiling characterizes diverse chondrocyte subtypes and nucleus pulposus progenitors in human intervertebral discs. Bone Res. Aug 16 2021;9(1):37. doi:10.1038/s41413-021-00163-z

12. Wang J, Huang Y, Huang L, et al. Novel biomarkers of intervertebral disc cells and evidence of stem cells in the intervertebral disc. Osteoarthritis and cartilage. 2021/03/01/ 2021;29(3):389–401. doi:10.1016/j.joca.2020.12.005

13. Panebianco CJ, Dave A, Charytonowicz D, Sebra R, Iatridis JC. Single-cell RNA-sequencing atlas of bovine caudal intervertebral discs: Discovery of heterogeneous cell populations with distinct roles in homeostasis. FASEB journal : official publication of the Federation of American Societies for Experimental Biology. Nov 2021;35(11):e21919. doi:10.1096/fj.202101149R

14. Calió M, Gantenbein B, Egli M, Poveda L, Ille F. The Cellular Composition of Bovine Coccygeal Intervertebral Discs: A Comprehensive Single-Cell RNAseq Analysis. Int J Mol Sci. May 6 2021;22(9)doi:10.3390/ijms22094917

15. Kuchynsky K, Stevens P, Hite A, et al. Transcriptional profiling of human cartilage endplate cells identifies novel genes and cell clusters underlying degenerated and non-degenerated phenotypes. Arthritis Res Ther. Jan 3 2024;26(1):12. doi:10.1186/s13075-023-03220-6

16. Risbud MV, Schoepflin ZR, Mwale F, et al. Defining the phenotype of young healthy nucleus pulposus cells: recommendations of the Spine Research Interest Group at the 2014 annual ORS meeting. J Orthop Res. Mar 2015;33(3):283–93. doi:10.1002/jor.22789

17. Melrose J, Smith SM, Fuller ES, et al. Biglycan and fibromodulin fragmentation correlates with temporal and spatial annular remodelling in experimentally injured ovine intervertebral discs. European spine journal: official publication of the European Spine Society, the European Spinal Deformity Society, and the European Section of the Cervical Spine Research Society. Dec 2007;16(12):2193–205. doi:10.1007/s00586-007-0497-5

18. Rajasekaran S, Tangavel C, Djuric N, et al. Part 1: profiling extra cellular matrix core proteome of human fetal nucleus pulposus in search for regenerative targets. Scientific reports. Sep 24 2020;10(1):15684. doi:10.1038/s41598-020-72859-x

19. Lin W, Chen H, Chen X, Guo C. The Roles of Neutrophil-Derived Myeloperoxidase (MPO) in Diseases: The New Progress. Antioxidants. 2024;13(1):132.

20. Breslin WL, Strohacker K, Carpenter KC, Haviland DL, McFarlin BK. Mouse blood monocytes: Standardizing their identification and analysis using CD115. Journal of Immunological Methods. 2013/04/30/ 2013;390(1):1–8. doi:10.1016/j.jim.2011.03.005

21. Austyn JM, Gordon S. F4/80, a monoclonal antibody directed specifically against the mouse macrophage. European Journal of Immunology. 1981;11(10):805–815. doi:10.1002/eji.1830111013

22. De Buck M, Gouwy M, Wang JM, et al. Structure and Expression of Different Serum Amyloid A (SAA) Variants and their Concentration-Dependent Functions During Host Insults. Curr Med Chem. 2016;23(17):1725–55. doi:10.2174/0929867323666160418114600

23. Worthley DL, Churchill M, Compton JT, et al. Gremlin 1 identifies a skeletal stem cell with bone, cartilage, and reticular stromal potential. Cell. Jan 15 2015;160(1-2):269–84. doi:10.1016/j.cell.2014.11.042

24. Bradley JE, Ramirez G, Hagood JS. Roles and regulation of Thy-1, a context-dependent modulator of cell phenotype. Biofactors. May-Jun 2009;35(3):258–65. doi:10.1002/biof.41

25. Goncharov NV, Popova PI, Avdonin PP, et al. Markers of Endothelial Cells in Normal and Pathological Conditions. Biochem (Mosc) Suppl Ser A Membr Cell Biol. 2020;14(3):167–183. doi:10.1134/s1990747819030140

26. Sawant KV, Poluri KM, Dutta AK, et al. Chemokine CXCL1 mediated neutrophil recruitment: Role of glycosaminoglycan interactions. Scientific reports. 2016/09/14 2016;6(1):33123. doi:10.1038/srep33123

27. Schall TJ, Bacon K, Toy KJ, Goeddel DV. Selective attraction of monocytes and T lymphocytes of the memory phenotype by cytokine RANTES. Nature. Oct 18 1990;347(6294):669–71. doi:10.1038/347669a0

28. Trapnell C, Cacchiarelli D, Grimsby J, et al. The dynamics and regulators of cell fate decisions are revealed by pseudotemporal ordering of single cells. Nature biotechnology. Apr 2014;32(4):381–386. doi:10.1038/nbt.2859

29. Alkhatib B, Ban GI, Williams S, Serra R. IVD Development: Nucleus pulposus development and sclerotome specification. Curr Mol Biol Rep. Sep 2018;4(3):132–141. doi:10.1007/s40610-018-0100-3

30. Mwale F, Roughley P, Antoniou J. Distinction between the extracellular matrix of the nucleus pulposus and hyaline cartilage: a requisite for tissue engineering of intervertebral disc. European cells & materials. Dec 15 2004;8:58–63; discussion 63-4. doi:10.22203/ecm.v008a06

31. Wang F, Cai F, Shi R, Wang XH, Wu XT. Aging and age related stresses: a senescence mechanism of intervertebral disc degeneration. Osteoarthritis and cartilage. Mar 2016;24(3):398–408. doi:10.1016/j.joca.2015.09.019

32. Kraus P, Lufkin T. Implications for a Stem Cell Regenerative Medicine Based Approach to Human Intervertebral Disk Degeneration. Front Cell Dev Biol. 2017;5:17. doi:10.3389/fcell.2017.00017

33. Moriguchi Y, Alimi M, Khair T, et al. Biological Treatment Approaches for Degenerative Disk Disease: A Literature Review of In Vivo Animal and Clinical Data. Global Spine J. Aug 2016;6(5):497–518. doi:10.1055/s-0036-1571955

34. Vadalà G, Ambrosio L, Russo F, Papalia R, Denaro V. Stem Cells and Intervertebral Disc Regeneration Overview-What They Can and Can’t Do. Int J Spine Surg. Apr 2021;15(1):40–53. doi:10.14444/8054

35. Walk R, Broz K, Jing L, et al. The progression of neurovascular features and chemokine signatures of the intervertebral disc with degeneration. bioRxiv. Jul 31 2024;doi:10.1101/2024.07.12.603182

36. Gelfand MV, Hagan N, Tata A, et al. Neuropilin-1 functions as a VEGFR2 co-receptor to guide developmental angiogenesis independent of ligand binding. eLife. Sep 22 2014;3:e03720. doi:10.7554/eLife.03720

37. Lee YJ, Lee HJ, Choi SH, et al. Soluble HSPB1 regulates VEGF-mediated angiogenesis through their direct interaction. Angiogenesis. Jun 2012;15(2):229–42. doi:10.1007/s10456-012-9255-3

38. Santambrogio L, Berendam SJ, Engelhard VH. The Antigen Processing and Presentation Machinery in Lymphatic Endothelial Cells. Front Immunol. 2019;10:1033. doi:10.3389/fimmu.2019.01033

39. Chen HY, Fermin A, Vardhana S, et al. Galectin-3 negatively regulates TCR-mediated CD4+ T-cell activation at the immunological synapse. Proceedings of the National Academy of Sciences of the United States of America. Aug 25 2009;106(34):14496–501. doi:10.1073/pnas.0903497106

40. Kano S, Sato K, Morishita Y, et al. The contribution of transcription factor IRF1 to the interferon-gamma-interleukin 12 signaling axis and TH1 versus TH-17 differentiation of CD4+ T cells. Nat Immunol. Jan 2008;9(1):34–41. doi:10.1038/ni1538

41. Bui TM, Wiesolek HL, Sumagin R. ICAM-1: A master regulator of cellular responses in inflammation, injury resolution, and tumorigenesis. J Leukoc Biol. Sep 2020;108(3):787–799. doi:10.1002/jlb.2mr0220-549r

42. Gschwandtner M, Derler R, Midwood KS. More Than Just Attractive: How CCL2 Influences Myeloid Cell Behavior Beyond Chemotaxis. Front Immunol. 2019;10:2759. doi:10.3389/fimmu.2019.02759

43. Rodrigues-Pinto R, Richardson SM, Hoyland JA. Identification of novel nucleus pulposus markers: Interspecies variations and implications for cell-based therapiesfor intervertebral disc degeneration. Bone Joint Res. 2013;2(8):169–78. doi:10.1302/2046-3758.28.2000184

44. Sun H, Wang H, Zhang W, Mao H, Li B. Single-cell RNA sequencing reveals resident progenitor and vascularization-associated cell subpopulations in rat annulus fibrosus. J Orthop Translat. Jan 2023;38:256–267. doi:10.1016/j.jot.2022.11.004

45. Sommerkamp P, Renders S, Ladel L, et al. The long non-coding RNA Meg3 is dispensable for hematopoietic stem cells. Scientific reports. Feb 14 2019;9(1):2110. doi:10.1038/s41598-019-38605-8

46. Wu S, Cheng Z, Peng Y, Cao Y, He Z. GPx3 knockdown inhibits the proliferation and DNA synthesis and enhances the early apoptosis of human spermatogonial stem cells via mediating CXCL10 and cyclin B1. Original Research. Frontiers in Cell and Developmental Biology. 2023-July-07 2023;11 doi:10.3389/fcell.2023.1213684

47. Fallik N, Bar-Lavan Y, Greenshpan Y, et al. Neat1 in hematopoietic stem cells. Oncotarget. Dec 12 2017;8(65):109575–109586. doi:10.18632/oncotarget.22729

48. Sekine K, Tsuzuki S, Yasui R, et al. Robust detection of undifferentiated iPSC among differentiated cells. Scientific reports. Jun 24 2020;10(1):10293. doi:10.1038/s41598-020-66845-6

49. Yu R, Han H, Chu S, et al. CUL4B orchestrates mesenchymal stem cell commitment by epigenetically repressing KLF4 and C/EBPδ. Bone Research. 2023/06/02 2023;11(1):29. doi:10.1038/s41413-023-00263-y

50. Surmann-Schmitt C, Dietz U, Kireva T, et al. Ucma, a novel secreted cartilage-specific protein with implications in osteogenesis. The Journal of biological chemistry. Mar 14 2008;283(11):7082–93. doi:10.1074/jbc.M702792200

51. Clayton SW, Walk RE, Mpofu L, Easson GWD, Tang SY. Sex-specific divergences in the types and timing of infiltrating immune cells during the intervertebral disc acute injury response are associated with reduced degenerative changes. Osteoarthritis and cartilage. Oct 18 2024; doi:10.1016/j.joca.2024.10.002

52. Butler A, Hoffman P, Smibert P, Papalexi E, Satija R. Integrating single-cell transcriptomic data across different conditions, technologies, and species. Nature biotechnology. Jun 2018;36(5):411–420. doi:10.1038/nbt.4096

