## Supplementary material for "Single cell RNA sequencing reveals shifts in cell maturity and function of endogenous and infiltrating cell types in response to acute intervertebral disc injury": {Supplemental Figures}

**Supplement:**

**Table S1**: Full List of Upregulated and Downregulated DEGS

**Table S2**: Panther Gene Ontology Terms Associated with Upregulated DEGs

**Table S3**: Panther Gene Ontology Terms Associated with Downregulated DEGs

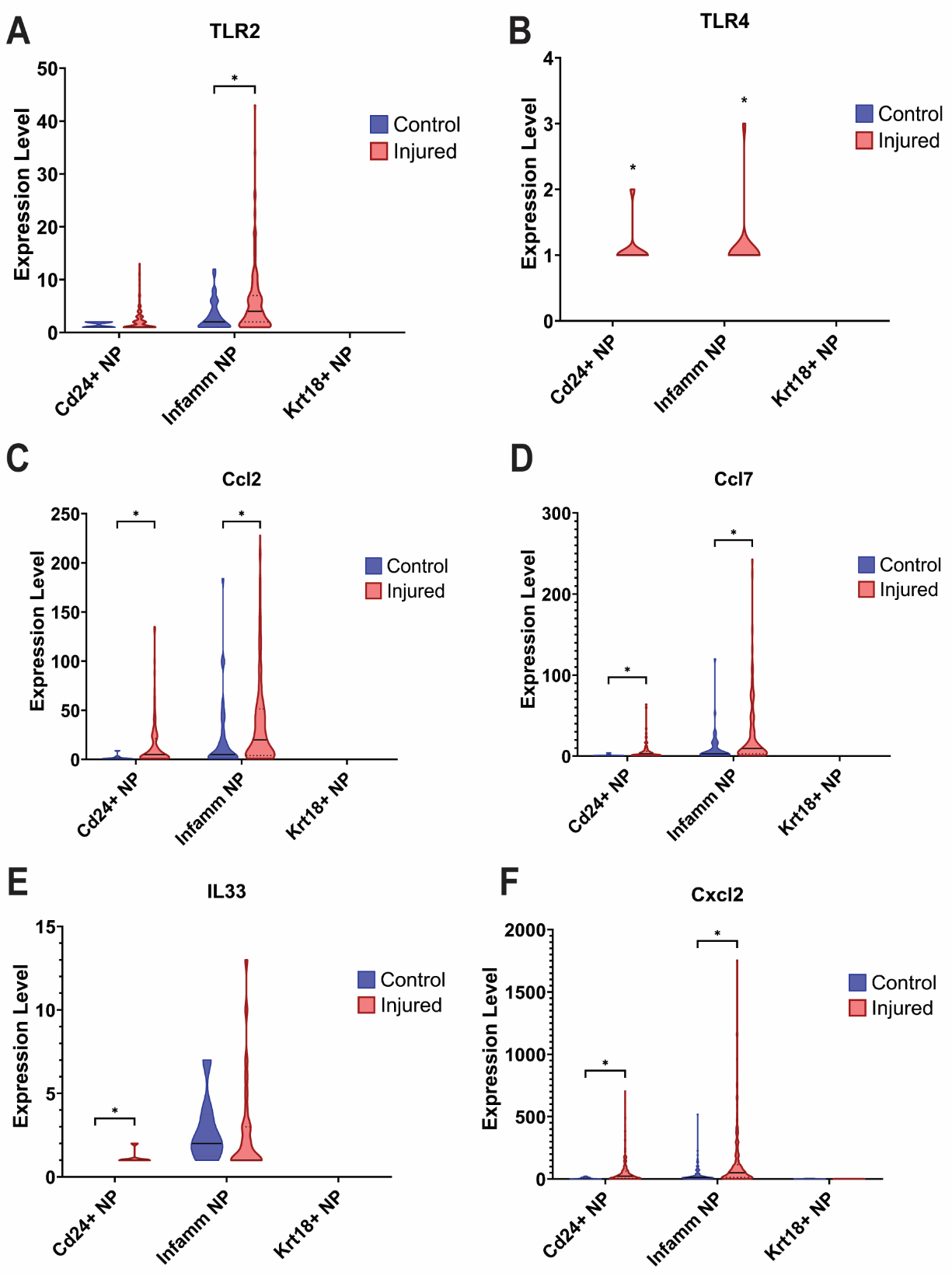

**Figure S1: The Inflammatory NP-Like cluster has higher expression levels of pro-inflammatory genes than the other NP clusters.** The Inflammatory NP-Like cluster has higher expression of proinflammatory Toll Like Receptors (A) *TLR2* and (B) *TLR4*, c, and cytokines (C) *Ccl2* (D) *Ccl7*, (E) *IL33*, and (F) *Cxcl2*

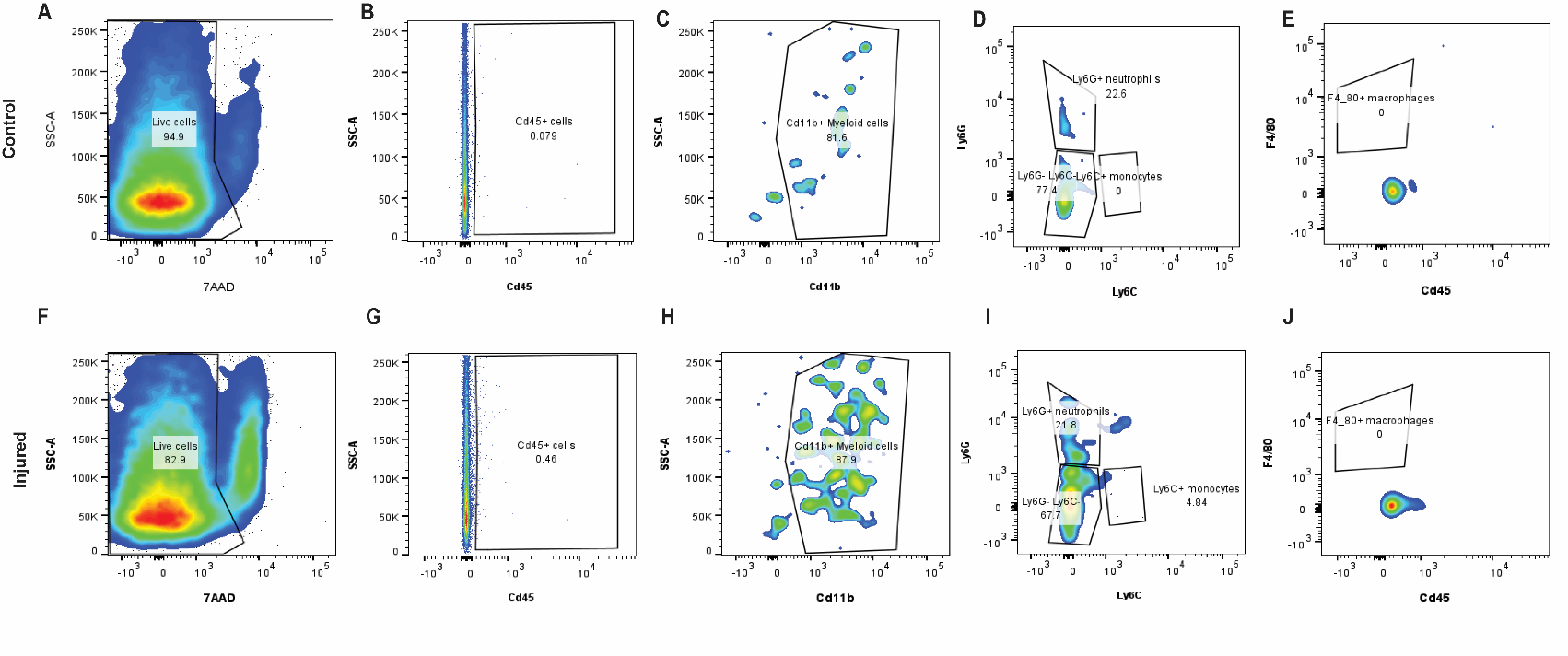
**Figure S2: Flow cytometry identified similar myeloid cells a those identified by scRNASeq at 7 dpi.** After selection of only the viable cells from (A) Control and (F) Injured IVDs, we identified an increase in Cd45^+^ immune cells with injury (B,G) and in the percentage of (C,H) Cd11b^+^ myeloid cells. (D,I) There are also similar percentages in neutrophils and an increase in monocytes with injury.

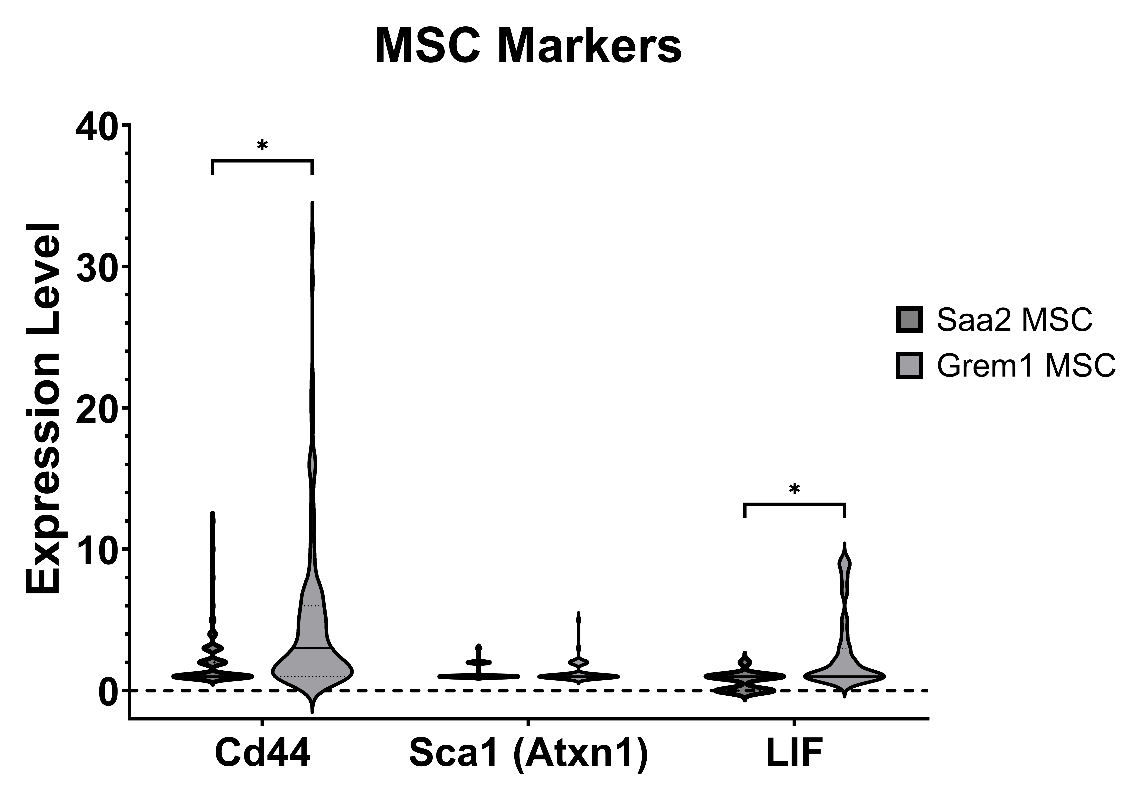

**Figure S3: Proliferative and stem cell marker expression from the MSC clusters.** The Saa2 MSC and Grem1 MSC clusters highly expressed stem cell markers: *Cd44*, *Sca1*, and *mKi67.* The Grem1 MSC cluster had higher expression levels of *Cd44* and *LIF* than Saa2 MSC cells.

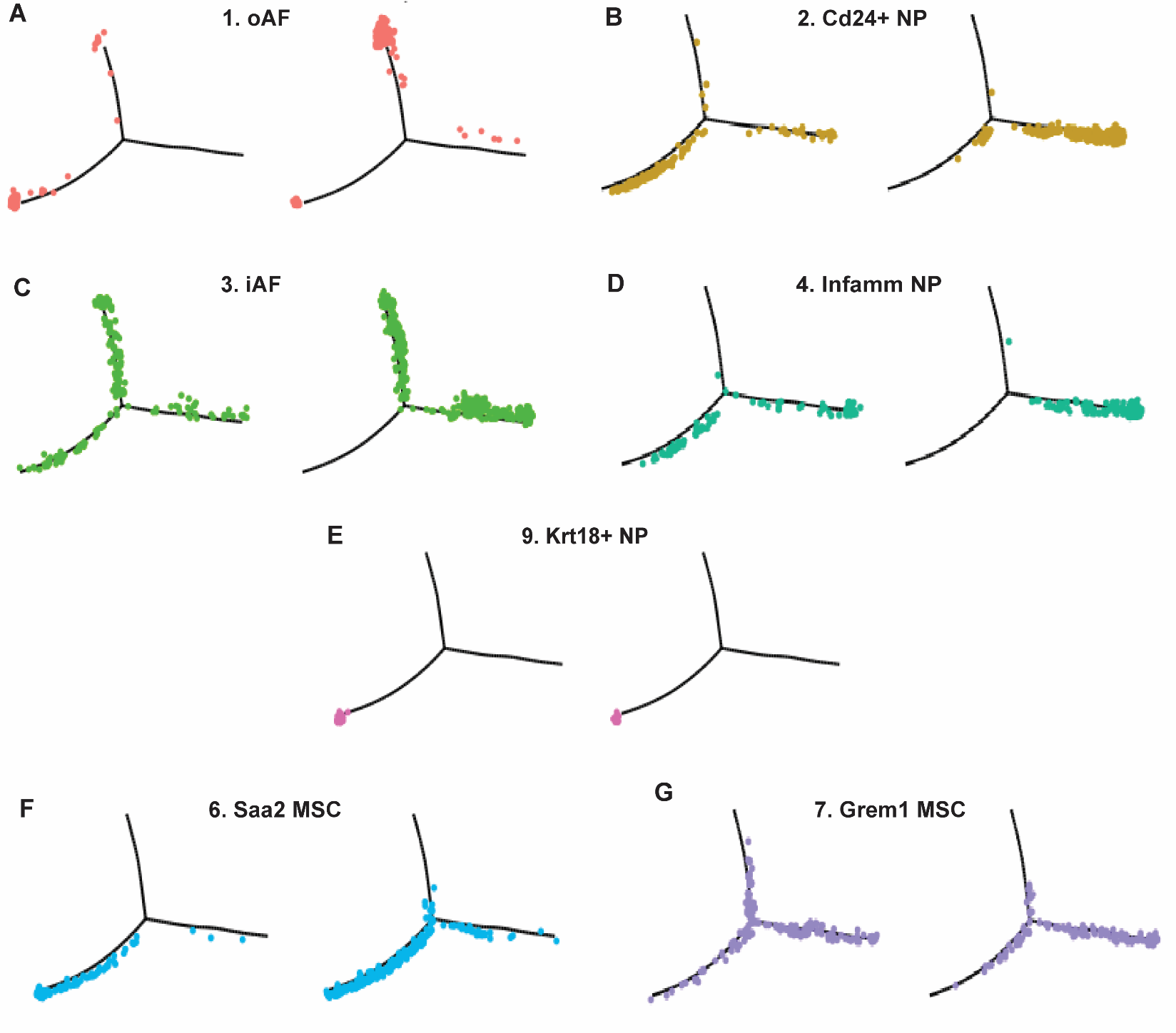

**Figure S4: Pseudotime trajectory analysis of each IVD and MSC cluster.** IVD tissue clusters and MSCs have distinct patterns of localization along the pseudotime trajectory tree that permits the identification and naming of the branches. (A) oAF cells majorly localize to the top branch with and without injury. (B) Cd24^+^ NP cells majorly localize to the bottom branch with and without injury, while (C) iAF cells localize to both branches. (D) Inflamm NP cells localize to the bottom branch and (E) Krt18^+^ NP cells remain in the most undifferentiated pseudotime position even with injury. Both (F) Saa2 MSCs retain some cells in the less differentiated pseudotime positions, and some cells localize to both branches with injury. (G) Grem1 MSCs have an increased number of cells that localize preferentially on the bottom branch and less cells in the undifferentiated pseudotime positions.

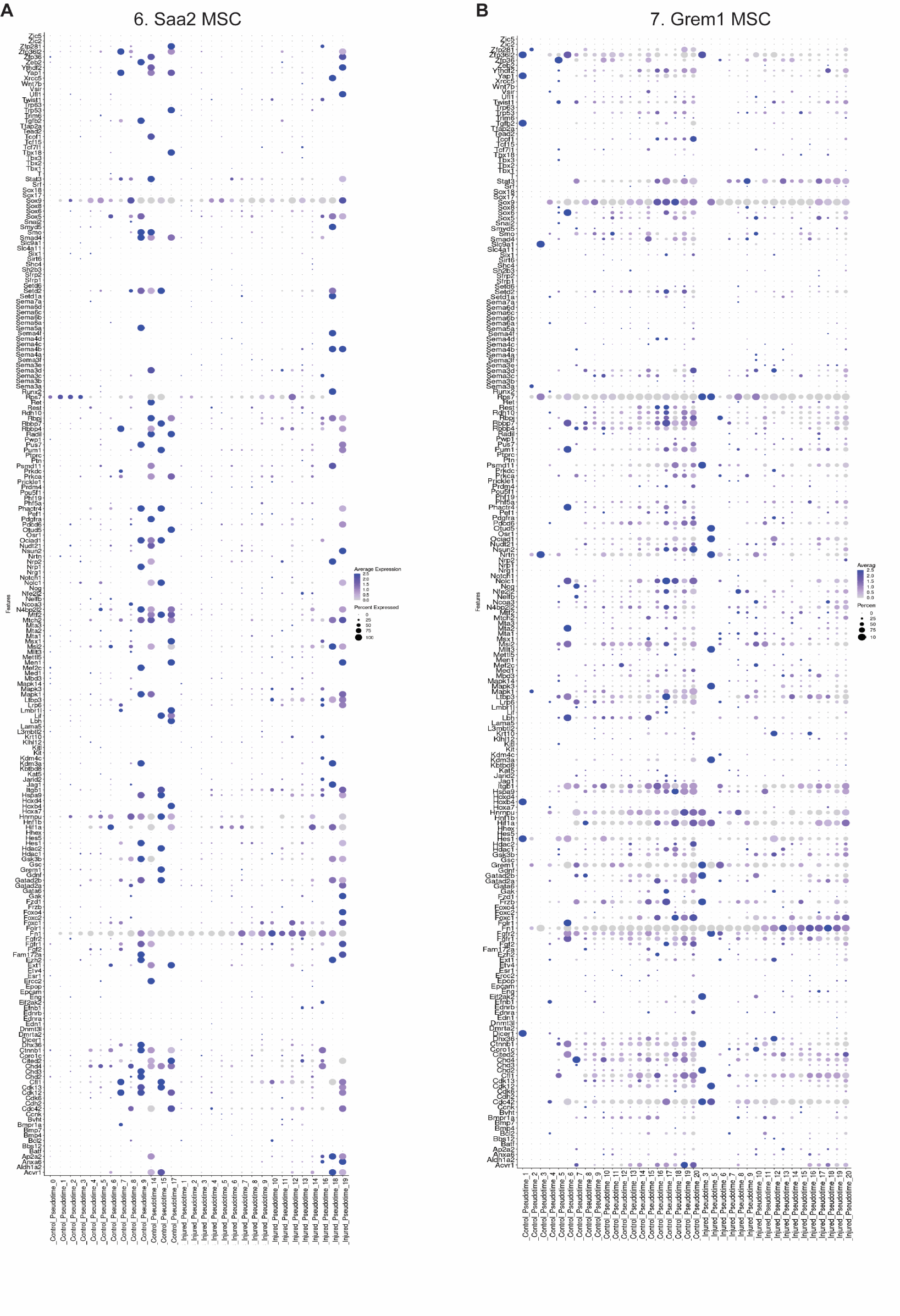

**Figure S5. Saa2 and Grem1 MSC Clusters Become Less Stem Cell Like Due to IVD Injury.** Both (A) Saa2 MSCs and (B) Grem1 MSCs have reduced gene expression levels and a reduction in the number of cells expressing general stem cell markers in the Injured cell pseudotime points.
